# Reward learning and working memory: effects of massed versus spaced training and post-learning delay period

**DOI:** 10.1101/2020.03.19.997098

**Authors:** G. Elliott Wimmer, Russell A. Poldrack

**Author notes:** Corresponding author*: G. Elliott Wimmer, Max Planck UCL Centre for Computational Psychiatry and Ageing, Wellcome Trust Centre for NeuroImaging, 10-12 Russell Square, University College London.

## Abstract

Neuroscience research has illuminated the mechanisms supporting learning from reward feedback, demonstrating a critical role for the striatum and midbrain dopamine system. However, in humans, short-term working memory that is dependent on frontal and parietal cortices can also play an important role, particularly in commonly-used paradigms in which learning is relatively condensed in time. Given the growing use of reward-based learning tasks in translational studies in computational psychiatry, it is important to understand the degree of working memory contributions and whether gradual learning mechanisms can be better isolated. In our experiments, we manipulated the spacing between repetitions along with a post-learning delay preceding a test phase. We found that learning was slower for stimuli repeated after a long delay (spaced-trained) compared to those repeated immediately (massed-trained), likely reflecting the remaining contribution of feedback learning mechanisms when working memory is not available. Brief interruptions of massed learning led to drops in subsequent performance. Further, individual differences in working memory capacity positively correlated with massed learning performance. Critically, after a delay period but not immediately, relative preferences decayed in the massed condition and increased in the spaced condition. Overall, our results provide additional support for a large role of working memory in reward-based learning in temporally condensed designs. We suggest that spacing training within or between sessions is a promising approach to better isolate and understand mechanisms supporting gradual reward-based learning, with particular importance for understanding potential learning dysfunctions in addiction and psychiatric disorders.

## Introduction

The accumulation of rewarding and aversive experiences exerts a strong influence on decision making. When making a choice between an apple and a banana, for example, an experienced decision-maker relies on values shaped by many experiences spread across years. In general, repeated experiences of stimulus- and action-reward associations are often separated by minutes, hours, days, or even longer, and recent research has shown that spaced training leads to value associations that are resistant to forgetting, similar to habits (Kim et al., 2015; Wimmer et al., 2018; van de Vijver and Ligneul, 2019). However, there is a striking difference between the proposed slow, gradual learning mechanism that supports the learning of habitual stimulus- and action-value associations (Yin and Knowlton, 2006) and actual experimental designs that are commonly used to study this learning mechanism in humans. Such “massed” designs feature closely-spaced repetitions and often feature rapidly shifting values (e.g. Daw et al., 2011; Wimmer et al., 2012; Wimmer et al., 2014).

Critically, recent research has begun to illuminate how performance in dominant massed reward learning paradigms is also supported by working memory processes (Collins and Frank, 2012; van de Vijver et al., 2015; Wimmer et al., 2018; van de Vijver and Ligneul, 2019). Working memory can maintain information – such as the identity of the best stimulus or the best response to a stimulus – in the face of interference, but has a limited capacity and a limited ability to store information over longer time periods (Ma et al., 2014; D’Esposito and Postle, 2015). Reward learning paradigms are increasingly being utilized in the growing field of computational psychiatry to study potential learning dysfunctions in mood and psychiatric disorders as well as addiction (Maia and Frank, 2011; Montague et al., 2012; Huys et al., 2016; Moutoussis et al., 2016). Performance differences in learning tasks between groups or across populations are often presumed to arise from differences in gradual striatal learning mechanisms. However, as a demonstration of potential violation of this assumption, it has been reliably shown that apparent deficits in reward-based learning in patients with schizophrenia are better accounted for by a deficit in working memory (Collins et al., 2014; Collins et al., 2017). Thus, it is important to further determine the contributions of working memory to ongoing reward-based learning, in order to understand how to better isolate the learning processes of interest.

While understanding interactions between massed training and working memory has been the focus of a number of recent studies (Collins et al., 2017; Collins, 2018; Collins and Frank, 2018), the inverse of massed training – spaced training – has been relatively under-explored in humans (Wimmer et al., 2018; van de Vijver and Ligneul, 2019). During learning, spacing between learning events has been associated with lower ongoing performance (Schmidt and Bjork, 1992; Taylor and Rohrer, 2010; Soderstrom and Bjork, 2015; van de Vijver and Ligneul, 2019). Critically, however, for many domains including verbal memory, motor skill learning, and educational performance, spacing between learning events is well-known to lead to reduced forgetting on later tests (reported in Ebbinghaus, 1913; Lee and Genovese, 1988; Donovan and Radosevich, 1999; Janiszewski et al., 2003; Cepeda et al., 2006). In the case of stimulus-reward association learning, it is unknown whether testing after a brief awake rest can lead to similar performance improvements.

To better understand the mechanisms underlying reward-based learning, our experiments examined the effect of massed versus spaced repetitions on ongoing learning and later test accuracy. We developed a single-session experimental paradigm that could provide multiple measures of this relationship and simultaneously examined the effect of spacing on performance during and after learning. Abstract stimuli were probabilistically paired with rewards or losses that depended on the participant’s response (**Fig. 1c**). Within-participants, training for a given “massed-trained” stimulus was completed in less than a minute, while training for a given “spaced-trained” stimulus was spread across approximately 15 min (**Fig. 1b**). During learning, to keep performance below ceiling participants also engaged in a secondary task (Waldron and Ashby, 2001; Foerde et al., 2006; Otto et al., 2013a). Following learning, in an across-participants manipulation, participants engaged in a choice test phase either immediately (the No Delay group) or after approximately 15 minutes (the Delay group; **Fig. 1a**).

**Figure 1.**
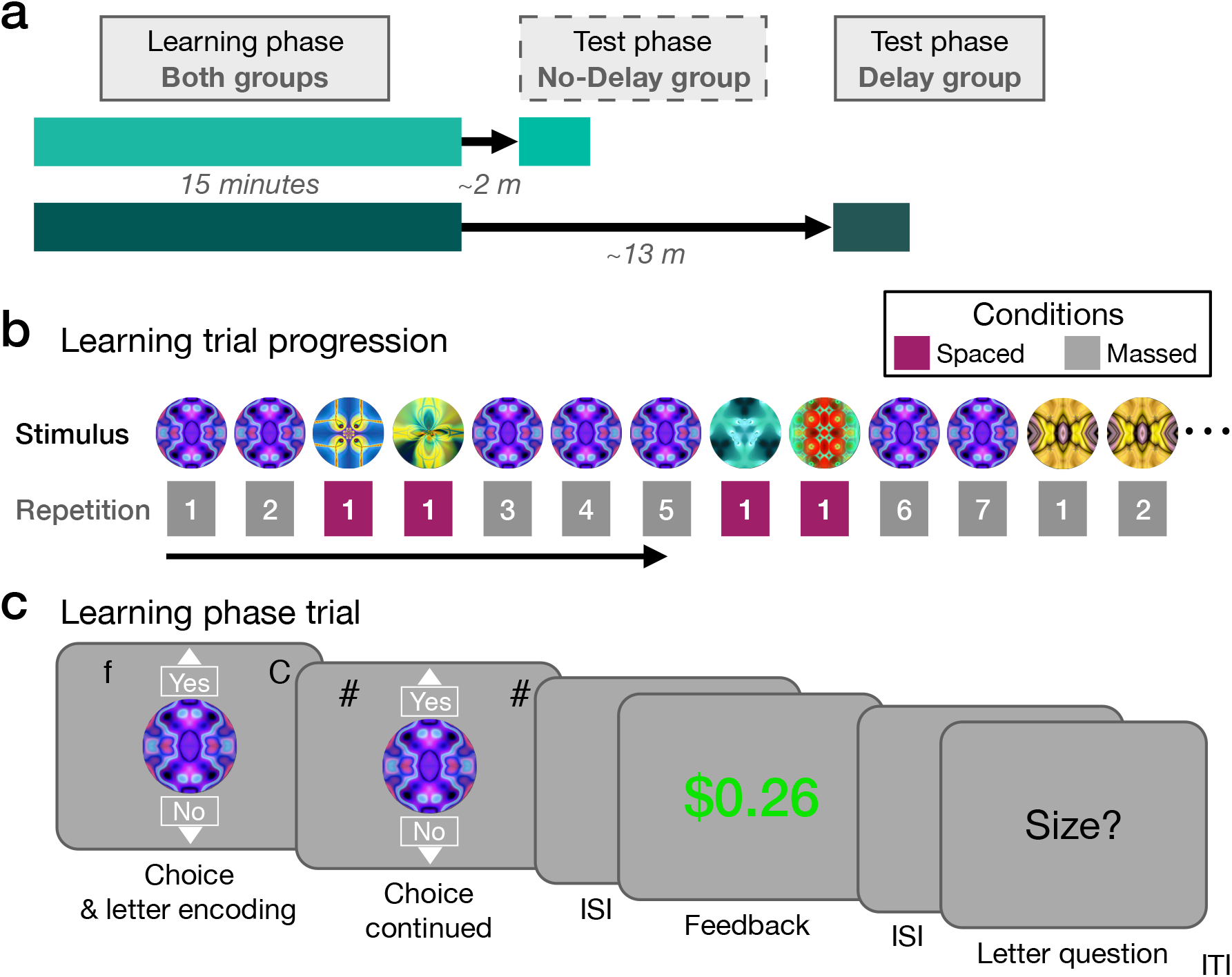
Experimental design and learning phase task. (**a**) Timeline of the learning phase and test phase (ratings, choices) for the No-Delay group and the Delay group. In the No-Delay group, the test phase began shortly after the completion of the learning phase (~2 min), while in the Delay group, the test phase began after a rest period (~13 min). (**b**) An example learning phase trial progression of massed- and spaced-trained stimuli, with the repetition number per stimulus noted below. Repetitions of massed-trained stimuli proceeded sequentially, with occasional interruptions by spaced-trained stimulus trials. (**c**) Reward learning phase. Participants made a ‘Yes’ or ‘No’ response to the abstract circle stimulus. In a secondary letter task, designed to partially occupy working memory and ensure below-ceiling learning performance, at the choice period, participants also encoded 2 letters. Following reward feedback, participants responded to a question about the letters.

We predicted that 1) Spaced training would lead to lower performance during ongoing learning; 2) Brief interruptions to massed training would be immediately followed by performance decreases; 3) Working memory capacity would positively relate to massed but not spaced performance during learning; and 4) Learned preferences for spaced-trained stimuli would show an increase relative to massed-trained stimuli following a rest period.

## Methods

### Participants

Participants were recruited via advertising on the Stanford Psychology paid participant pool web portal. Informed consent was obtained in a manner approved by the Stanford University Institutional Review Board. Target sample size was based on a related study of spacing and reward-based learning (Wimmer et al., 2018). In the Delay group, from an initial number of 35 participants, 8 were excluded based on learning performance according to the methodology described below, leaving a total of 27 participants (13 female; mean age = 22.2, range: 19-29). In the No-Delay group, from an initial number of 34 participants, 7 were excluded based on performance, leaving a total of 27 participants (15 female; mean age = 21.7; range: 18-30). Participants were paid $10/hour for the approximately 75-minute experiment, plus monetary rewards from the learning phase and choice test phase, leading to total compensation of approximately $20.

Participants were excluded based on learning phase performance measures unrelated to our hypotheses of interest. We did not use the rating or choice test phases for exclusion. First, we excluded participants with mean massed learning performance on the last two repetitions of ≤ 59% (1 exclusion in the Delay group; 5 in the No-Delay group). Near-chance learning performance in the massed condition is likely related to poor overall attention to the reward learning task. Second, we excluded participants with an extreme Yes/No response bias for the spaced stimuli in the second half of repetitions (5-7), where ≥ 90% of responses were of the same option across all spaced stimuli (3 exclusions in the Delay group; 1 in the No-Delay group). Strong response bias indicates an overreliance on a fixed strategy for the spaced stimulus trials, which likely interferes with non-strategic stimulus-specific learning mechanisms. Third, we excluded participants with poor performance on the letter judgment secondary task (≤ 59% accuracy; 4 exclusions in the Delay group; 2 in the No-Delay group, where 1 also met the response bias exclusion criteria). Near-chance performance indicates a lack of attention to the secondary task. As performance in the dual-task was lower than expected overall, we used an exclusion threshold of ≤ 59% accuracy. Raising the performance exclusion threshold for the letter task did not qualitatively affect the results.

### Experimental Design

The reward learning task included three phases: a learning phase, a rating test phase, and a choice test phase. The No-Delay and Delay groups differed only in a potential delay between the end of the learning phase and the beginning of the test phase. In the Delay group, an approximately 13 min break was inserted between the last learning phase trial and the first rating phase trial. In the No-delay group, an approximately 2 min break was inserted between the last learning trial and the first rating phase trial. The delay or “rest” period in the Delay group was intended 1) to provide a pause during which potential forgetting as well as offline replay processes might differentially affect spaced-trained versus massed-trained value associations (Gershman et al., 2014), and 2) to better equalize the relative delays between initial learning and subsequent testing for the massed stimuli, as exposure to the final massed stimulus was temporally quite proximal to the start of the ratings test phase in the No-Delay group. During the delay, participants were told that they were to take a break and could occupy themselves by leaving the testing room, interacting with their mobile devices, or browsing the internet on the testing room computer.

The learning phase was similar to a simple reward-based learning task we have used previously (Gerraty et al., 2014; Wimmer et al., 2018) (**Fig. 1c**). Abstract circle stimuli were associated with either potential reward or potential loss, depending on the participant’s response. Reward or loss associations were only revealed through trial-and-error learning. The goal was to learn the best response for each stimulus (arbitrarily labeled “Yes” and “No”) across stimulus repetitions, in order to accumulate as large a bonus as possible. For reward-associated stimuli, the optimal response was to select “Yes” in order to win a potential gain most of the time (mean +$0.25, in green font; 86% probability; versus a 14% probability of a small loss; mean −$0.05, in grey); the outcome probabilities were inverted for a “No” response. For loss-associated stimuli, the optimal response was to select “No” in order to achieve a neutral outcome most of the time (mean +$0.00, in grey; 86% probability; versus a 14% probability of a loss; mean −$0.25, in red); the outcome probabilities were inverted for a “Yes” response. To increase engagement, displayed reward amounts were jittered by adding a uniform distribution of ±5 cents around the mean. Concurrent with the reward learning task, to add a load to working memory, participants also engaged in a secondary letter task (**Fig. 1c**).

Stimuli were either “massed” or “spaced”, distinguished by the distance between stimulus repetitions. Each stimulus was repeated seven times. All repetitions of an individual massed stimulus were in sequence, with occasional interruptions by 1-4 spaced stimulus trials (mean 2.73), with all repetitions for a given massed stimulus spread across ~1 min. The repetitions of each spaced stimulus were spread across the full learning phase of ~15 min. Thus, spacing was manipulated within the same session, in contrast to a multi-session spacing paradigm we reported recently (Wimmer and Poldrack, 2018); see also a very similar approach by van de Vijver and Ligneul (2019). For the spaced stimuli, the six different stimuli were each presented a single time in a pseudo-random order before continuing with the next repetition, with no direct repetitions of the same stimulus. The average total trial separation (including massed and spaced stimuli) between repetitions of spaced-trained stimuli was 14.2 trials, with a minimum of 5 and a maximum of 28. For spaced stimuli, an interruption by a spaced trial (or trials) began with an approximately equal number of reward and loss stimuli. Five of the seven transitions between different massed stimuli were marked by the presence of one or more spaced trials. For massed stimuli, no more than two stimuli associated with reward (or loss) followed in sequence. Stimulus assignment to the massed and spaced conditions were counterbalanced across participants.

In order to ensure that repetitions of massed stimuli were sufficiently close together in the learning phase, minimizing the number and duration of interruptions by spaced stimuli, it was necessary to include more massed stimuli than spaced stimuli. Thus, the experiment included 8 massed-trained stimuli and 6 spaced-trained stimuli (yielding a total of 98 learning trials). It is unlikely that the different numbers of stimuli in the two conditions affected our results. We predicted that learning in the massed condition would be primarily supported by working memory while learning in the spaced condition would be primarily supported by gradual stimulus-response learning. Including a larger number of stimuli overall (or just in the massed condition) would be expected to decrease relative performance in the massed condition by taxing working memory load. Consequently, this would, if anything, decrease our ability to detect the predicted differences between conditions, specifically, relatively higher massed learning performance at the end of training and a relative decrease in preference strength for massed stimuli at a delayed test. Further, if working memory load was increased by the number of massed stimuli, this would increase the relative contribution of a gradual stimulus-response learning mechanism in the massed condition (Collins, 2018), again decreasing our ability to detect any differences between conditions. Alternatively, if learning in both conditions was supported by gradual stimulus-response (model-free) learning such as that associated with the striatum, storing response associations has a minimal memory cost and so the number of stimuli per condition is negligible (Collins and Frank, 2012). Conversely, if learning in both conditions was supported by short-term working memory, each stimulus, independent of spacing condition, would add an additional element to be stored, but this would not affect one condition more than the other.

To detail the events on a reward learning trial, a stimulus was first presented with the options “Yes” and “No” above and below the image, respectively (**Fig. 1c**). Above the circle, for the concurrent short-term memory task, two letters were presented on the left and right sides of the screen. The letters appeared on the screen for 0.30 s before being replaced by pound signs (“#”) for 0.20 s. The participants could make their Yes/No response to the circle stimulus at any time during the letter presentation phase or afterwards. Participants used the up and down arrow keys to make “Yes” and “No” responses, respectively, within the full 2 s choice period. After a response, the Yes/No options remained on the screen for the remainder of the period. A 1 s blank inter-stimulus interval followed. Reward feedback was then shown in text in the center of the screen for 1.5 s. If a response was not made in the choice period, participants were shown text stating “Too late or wrong key! - $0.50” in red font. A brief 0.25 s blank inter-stimulus interval followed. Next, either the question “Earlier?” or “Size?” was presented, indicating that the relevant question about the letters shown during choice would be about position in the alphabet or capitalization, respectively. Participants had 2 s to make a response, using the left and right arrow keys. A blank ISI of 0.50 s followed. If an incorrect response was made or if no response was recorded, “Incorrect!” appeared on the screen in red font for 0.75 s. If a correct response was made, a brief fixation of 0.25 s followed. Finally, an inter-trial-interval including a white fixation cross was presented for an average of 2 s (range: 0.50-3.25 s), followed by a trial-start indicator where the white fixation changed to black for 0.25 s.

We adapted our secondary task (which was concurrent with the learning phase) from previous work (Waldron and Ashby, 2001; Otto et al., 2013a), using letters instead of numbers to avoid interference with numerical reward feedback amounts. Letters were taken from the set of letters ‘a’ through ‘j’, excluding ‘i’. Letters on each trial were balanced such that the larger or earlier letter appeared approximately equally on the left and right side of the screen, leading to an approximately even distribution of the correct response to the left and right options. On ~85% of trials, the correct answers for the two potential probe questions were different; on the remaining trials, one letter was both earlier and a capital letter. Mid-way through the practice block and approximately every 25 trials thereafter, the computer displayed a warning if letter task performance fell below 66%.

The learning phase began with 12 practice trials, including one reward- and one loss-associated practice stimulus, during which the letter presentation period was increased by 0.50 s and the choice response period was increased by 2 s.

After the learning phase, in the Delay group, participants first had a break of ~13 min (mean 12.76 min, range 7.6-22.5 min). In the No-Delay group, the time between the last learning trial and the first rating trial was ~2 min (mean 2.04 min, range 1.4-5.0 min), allowing for the experimenter to return to the testing room and administer instructions. In the reward rating phase, participants saw each abstract circle stimulus and tried to recall whether that stimulus was associated with reward or loss. Below the stimulus, a rating scale appeared, anchored by “0% reward” on the left and “100% reward” on the right. Participants were instructed to try to remember the value of a stimulus, using their best guess or gut feeling. They were instructed that the endpoints of the scale represented complete confidence in their answer, while points closer to the middle indicated lower confidence. Participants indicated their response using a computer mouse (with no time limit), followed by a 3 s ITI. The phase began with a single practice trial followed by the massed and spaced stimuli in a pseudo-random order.

Next, participants completed the incentive-compatible choice test phase, our primary test measure. On each trial, one stimulus was presented on the left and an alternative stimulus was presented on the right, with left-right location randomized. Participants were instructed to choose the stimulus that they thought had been associated with reward over the stimulus they thought had been associated with loss. Further, participants were informed that they would not receive feedback, but that choices of the reward-associated stimulus would add to their monetary reward at the end of the experiment. Participants were instructed to use their best guess or gut feeling. Participants made their responses using a 4-point scale: “(1) Sure left, (2) Guess left, (3) Guess right, (4) Sure right” using the 1-4 keys. They were further instructed that the level of confidence of their answer did not affect the potential bonus for correct choices. A 3.5 s ITI followed the response. The phase began with a single practice trial using the two stimuli from the practice learning trials followed by the choice trials in pseudo-random order.

The primary choice trials contrasted a reward vs. a loss-associated stimulus, where both stimuli came from the spaced condition or both from the massed condition. The phase also contained a secondary kind of choice comparing a stimulus from the spaced to the massed condition, where both had the same value association (both reward or both loss). In the Delay group, 43 choices were presented. All of the potential combinations of the three reward versus three loss stimuli from the spaced condition were presented (resulting in 9 choices). A subset of the choices from the massed condition were presented (resulting in 10 choices; a subset was used in order to reduce potential fatigue). The remainder of the choices were across-condition choices. Choice order was pseudo-randomized. The order was set to that the first 8 choices were all primary within-condition choices while the last 12 choices were between-spacing condition choices. The choice test phase in the No-Delay group used the same set of choices as the Delay group, including the full set of within-condition massed choices, yielding 49 choice trials. After the choice test phase, participants completed a short written questionnaire.

In the No-Delay group, we additionally administered a working memory measure, the operations span task (O-SPAN) (Lewandowsky et al., 2010; Otto et al., 2013b). In the O-SPAN, participants made accuracy judgments about simple arithmetic equations (e.g. ‘2 + 2 = 5’). After a response, a to-be-encoded letter appeared (e.g. ‘B’), followed by the next equation. Arithmetic-letter sequences ranged in length from 4 to 8. At the completion of a sequence, participants were asked to type in the letters that they had seen in the original order, with no time limit. Each of the sequence lengths was repeated 3 times with different equations and letters in a pseudo-random order. In order to ensure that participants were fully practiced in the task before it began, the task was described in detailed instruction slides, followed by 5 practice trials. Scores were calculated by summing the number of letters in fully correct letter responses across all 15 trials (mean, 50.7; range, 19-83) (Otto et al., 2013b; Wimmer et al., 2018). All participants maintained a level of correct arithmetic performance above 70%, with group mean performance of 94%.

### Behavioral analysis

Behavioral analyses were primarily conducted in Matlab 2018b (The MathWorks, Inc., Natick, MA). Interactions between delay group and spacing condition were examined via ANOVA, using the function anovan. Learning performance was quantified as percent correct choice (“Yes” for the reward-associated stimuli, and “No” for the loss-associated stimuli) and compared to chance using a t-test. To examine the effect of interspersed spaced trials on concurrent massed learning performance, the performance change from pre- to post-interruption was compared to a balanced control performance change across repetitions with no interruption. The control non-interruption performance measure was constructed by computing an average of trial-to-trial performance changes weighted by the actual number of times that massed learning was interrupted at a given learning repetition. In this way, the balanced control measure was used to compare interrupted versus non-interrupted performance changes from repetition 3 to 4 and repetition 4 to 5, with a smaller weight given to changes from rarer interruptions for repetitions 2 to 3, 5 to 6, and 6 to 7.

Test phase choices were averaged within each spacing condition. We also corrected choice accuracy by performance at the end of learning, using the last repetition from the learning phase. Note that while learning phase performance and choice accuracy use different behavioral measures, any interaction between change in performance and delay group only depends on relative differences in performance, independent of the different underlying measures. Test phase reward ratings were recorded on a graded scale from 0 to 100, where 50 indicated neutral. Per-participant regression models estimated the relationship between reward ratings (with one value per stimulus) and test phase choices. At the second level, coefficients were compared via ANOVA. In further analyses, ratings variability was computed as the mean of reward and loss stimulus ratings variability.

Learning phase multilevel regression analyses were conducted in R (https://www.r-project.org/). We used lme from the nlme package for linear regression and glmmTMB from the glmmTMB package for logistic regression. All predictors and interactions were included as random effects, following the ‘maximal’ approach (Barr et al., 2013). Correlations between random effects were included when convergence was achievable with this structure. The primary logistic regression learning model examined the relationship between group (No-Delay, Delay), spacing condition (massed, spaced), repetition (1 – 7), and all interaction effects on correct responses. A secondary analysis examined the effect of reward versus loss association on correct responses. Equivalent models examined letter task accuracy. All reported p-values are two-tailed.

We additionally examined correlations between working memory capacity (as measured with the O-SPAN) and behavioral learning performance. Individual differences in performance from the learning phase were based on behavior after sufficient task exposure (here, the second half of the learning phase) in order to prevent confounding factors such as initial task adjustment, attentional orienting, and task-set learning from contributing noise to any potential relationship, following previous procedures (Wimmer and Poldrack, 2018). Correlations were computed using Pearson’s correlation. Statistical comparison of the difference in working memory correlations with massed versus spaced learning performance was computed using Steiger’s test for differences in dependent correlations.

For all results of interest, we tested whether non-significant results were weaker than a moderate effect size using the Two One-Sided Test (TOST) procedure (Schuirmann, 1987; Lakens, 2017) as implemented in the TOSTER library in R (Lakens, 2017). We used bounds of Cohen’s *d* = 0.57, where power to detect an effect in the included group of n = 27 participants in either the No-Delay or Delay group is estimated to be 80%. For effects across all 54 participants, to achieve 80% power we used bounds of 0.40. For correlations, the *r*-value cutoff for 80% power was estimated to be 0.37.

## Results

### Learning

#### Effect of spacing on learning performance

The learning phase procedure was the same across the No-Delay and Delay groups, and we thus expected similar learning performance across delay groups.

Overall, performance increased across the learning phase, as demonstrated by an effect of stimulus repetition on performance (multilevel regression model *β* = 0.196 [0.165 0.228]; z = 12.35, p < 0.0001; similar effects were found in each group separately; **Table 1; Fig. 2a**). To confirm that learning was similar across delay groups, we tested for and found no significant three-way interaction between repetition, delay group, and spacing condition on learning performance (*β* = −0.010 −0.078 0.059]; z = −0.279, p = 0.781; TOST p = 0.005; indicating that we can rule of the possibility of a medium-sized effect or larger). We also found no significant interaction between repetition and delay group on learning performance (*β* = −0.026 [-0.057 0.004]; z = −1.645, p = 0.10; TOST p = 0.10; thus we cannot rule out the possibility of a medium-sized effect). Separately, in a secondary analysis, we also found that performance was higher for reward- versus loss-associated stimuli (*β* = 0.621 [0.425 0.955]; z = 5.103, p < 0.0001; group interaction p = 0.821).

**Table 1.** Mean learning phase performance separated by group (No-Delay, Delay) and condition (massed, spaced) for the last stimulus repetition in the reward learning task (top) and across the phase for the concurrent secondary letter task (bottom). CI = 95% confidence intervals.

| Learning | Condition | Final mean | CI |
| --- | --- | --- | --- |
| No-delay | massed | 82.0 | [76.9 87.1] |
|  | spaced | 57.2 | [50.1 64.2] |
| Delay | massed | 89.2 | [84.7 93.7] |
|  | spaced | 57.9 | [50.1 65.7] |

| Letter task | Condition | Mean | CI |
| --- | --- | --- | --- |
| No-delay | massed | 81.5 | [78.2 84.8] |
|  | spaced | 80.1 | [76.4 83.7] |
| Delay | massed | 74.9 | [71.3 78.6] |
|  | spaced | 76.0 | [71.8 80.3] |

**Figure 2.**
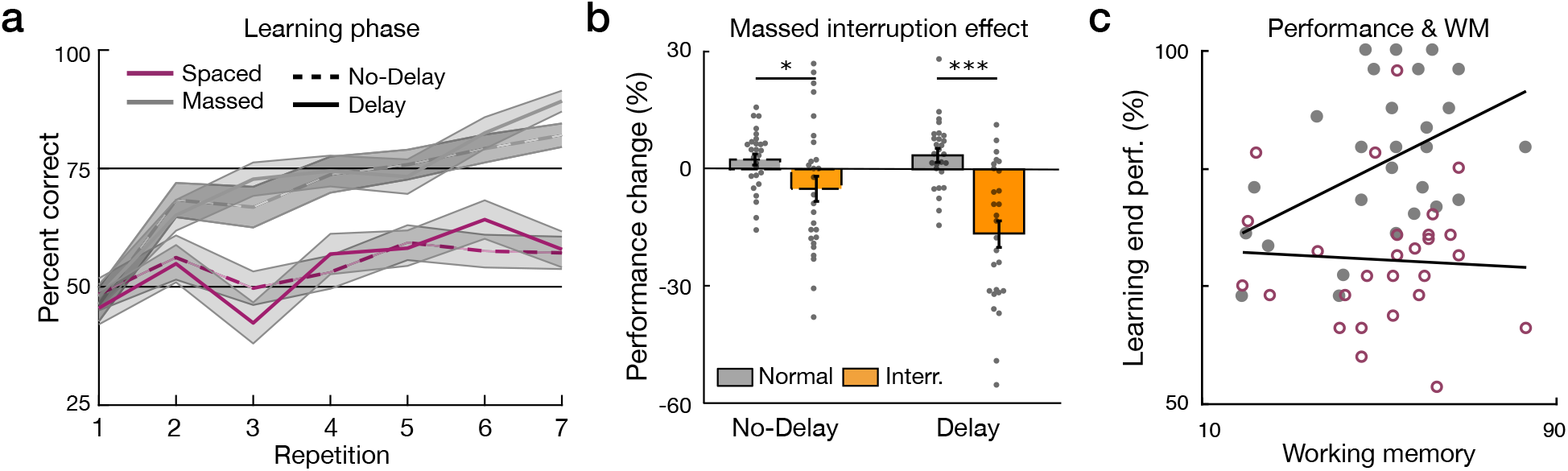
(**a**) Learning performance in the No-Delay group (dashed lines) and Delay group (solid lines) across the learning phase for massed-trained (grey) and spacedtrained (magenta) stimuli. Shaded error margins represent standard error of the mean (SEM). (**b**) During learning, interruption of massed training by occasional sets of spaced trials was followed by a relative decrease in post-interruption performance in both the No-Delay and Delay group (blue), relative to the normal learning-related increase in performance in a balanced no-interruption repetition measure (grey). Dots represent individual participants; outlines are dashed in the No-Delay group. (**c**) A working memory measure was collected in the No-Delay group (O-SPAN). Working memory capacity was positively correlated with massed but not spaced learning performance (difference p = 0.004; massed condition, filled grey circles; spaced condition, open magenta circles). (*p < 0.05; **p < 0.01; ***p < 0.001; for underlying data, see https://osf.io/x5u3n/, https://osf.io/vkj8n/, and https://osf.io/wj3va/)

We predicted that performance for the immediately-repeating massed-trained stimuli would be partially supported by working memory, while performance for spaced-trained stimuli would not benefit from this assistance. Consistent with this prediction, we found that performance for massed-trained stimuli was significantly higher than performance for spaced-trained stimuli overall (*β* = 0.837 [0.679 0.994]; z = 10.419, p < 0.0001). Further, the massed performance benefit was also reflected in a stronger effect of repetition in the massed versus spaced conditions (*β* = 0.200 [0.132 0.269]; z = 5.734, p < 0.0001).

Overall, learning performance in the spaced condition was quite low. As designed, the long spacing between stimulus repetitions minimized the contribution of short-term working memory. Further, we speculate that the low performance for spaced-trained stimuli could be due to the demanding secondary task adding noise to the choice process. An additional possibility is that by occupying attentional resources, the secondary task decreases the contribution of an additional learning mechanism, episodic memory, to association learning (Foerde et al., 2006; Wimmer and Buchel, 2016; Gershman and Daw, 2017).

#### Learning phase letter task performance

To keep massed performance below ceiling, participants engaged in a demanding secondary task which required them to remember letter identity and letter size for two letters over the course of each trial (e.g. ‘B’ and ‘e’; **Fig. 1c**). In both groups performance on the letter task was far above chance (p-values <0.0001). We found no effect of spacing and no interaction between spacing and delay group on letter task performance (p-values > 0.34; **Table 1**). However, we did find an effect of group (*β* = 0.165 [0.025 0.306]; z = 2.305, p = 0.021), such that performance in the No-Delay group was higher than in the Delay group.

Similar to reward learning performance, in a secondary analysis we found that performance on the letter task in both groups was higher for reward-associated stimuli than loss-associated stimuli (β = 0.435 [0.279 0.590]; z = 5.470, p < 0.0001; group interaction p = 0.219). The difference between letter task performance on reward-versus loss-associated stimulus was also greater in the massed- than spaced-trained condition (β = 0.623 [0.354 0.912]; z = 4.449, p < 0.0001; group interaction p = 0.213). The performance benefits in the learning task for reward-associated stimuli, as reported above,, may lead to the observed higher performance on reward trials in the secondary letter task due to decreased interference from learning-related processes.

#### Effect of interruptions on massed learning performance

In a second test of the prediction that performance on massed-trained stimuli would be assisted by short-term memory, we examined the effect of interruptions of massed stimulus repetitions by occasional interleaved spaced stimulus trials (**Fig. 1b**). Across the seven repetitions of a given massed stimulus, spaced stimulus trials were pseudo-randomly inserted (median 2 spaced trials per interruption, range 1-3). Supporting our prediction, across groups we found that the interruption of massed stimulus repetitions by spaced trials negatively affected post-interruption performance (post-pre interruption performance versus no-interruption control; F_(2,104)_ = 27.97, p <0.0001; η_p_^2^ = 0.212; **Fig. 2b**). While the learning phase was the same in both groups, we found a stronger interruption effect in the Delay group (F_(2,104)_ = 5.90, p = 0.017; η_p_^2^ = 0.054). Planned comparisons confirmed that the interruption effect was significant in both groups (No-Delay t_(26)_ = 2.318, 95% confidence interval (CI) [0.009 0.141]; p = 0.029; Delay t_(26)_ = 5.280, CI [0.123 0.280]; p < 0.001; **Fig. 2b**).

The effect of interruption was also reflected in reaction time, such that reaction time exhibited a transient increase post-interruption (F_(2,104)_ = 20.40, p <0.0001; η_p_^2^ = 0.164). We found no difference in the interruption effect on reaction time across delay groups (F_(2,104)_ = 0.32, p = 0.574; η_p_^2^ = 0.003). Planned comparisons confirmed that the interruption effect was present in both groups (No-Delay t_(26)_ = −3.015, CI [−0.121 −0.023]; p = 0.006; Delay t_(26)_ = −3.571, CI [−0.146 0.039]; p = 0.0014).

#### Massed learning performance and working memory

As a third test of the relationship between performance in the massed condition and working memory, we expected that overall massed condition learning performance would be related to individual differences in working memory capacity (Wimmer and Poldrack, 2018). In the No-Delay group, we collected a separate measure of working memory capacity (O-SPAN). We found that performance in the massed but not the spaced condition was significantly correlated with working memory (r = 0.56, p = 0.0024; spaced, r = −0.20, p > 0.31; **Fig. 2c**). Further, the performance-WM correlation in the massed condition was significantly stronger than that in the spaced condition (z = 2.86, p = 0.004). Thus, three separate behavioral measures support a role for short-term memory in rapid massed learning: First, learning for massed-trained stimuli was faster than learning for spaced-trained stimuli. Second, learning for massed-trained stimuli was negatively affected by interruptions by spaced stimulus trials. Third, massed learning performance was positively correlated with individual differences in working memory capacity.

### Choice test

#### Effect of spacing and delay on learning maintenance

After the learning phase, we tested participants’ memory for learned associations in incentive-compatible choices between reward- and loss-associated stimuli, where the Delay group experienced a rest preceding the test phase. Our analyses focus on choice data corrected for performance at the end of the learning phase. All results focused on the critical choices between stimuli within each spacing condition; for choices across spacing conditions see Supplementary Results.

First, to ensure that participants successfully chose reward-associated stimuli over loss-associated stimuli, we examined raw choice preference data prior to correcting for performance at the end of the learning phase. In the choice measure unadjusted for end-of-learning performance, we found no significant interaction between delay group and spacing condition in the uncorrected choice data (F_(2,104)_ = 2.54, p = 0.114; η_p_^2^ = 0.024; **Table 2**). In both the No-Delay and Delay groups, choice preference for reward- over loss-associated stimuli was greater than chance for both massed- and spaced-trained stimuli (p-values < 0.036, corrected for multiple comparisons; **Table 2; Figure 3a**). In planned comparisons, in the No-Delay group, we found that test phase accuracy was higher for massed- vs. spaced-trained stimuli (t_(26)_ = 2.81, CI [3.7 23.7]; p = 0.019; **Fig. 3a**), similar to performance at the end of the preceding learning phase. In contrast, in the Delay group we found that choice accuracy was numerically matched for massed- and spaced-trained stimuli (t_(26)_ = −0.13, CI [−11.6 10.3]; p = 0.90; TOST p = 0.004; **Fig. 3a**).

**Table 2.** Uncorrected choice test phase accuracy (top) and accuracy change from the end of learning to the choice test (bottom) by group (No-delay, Delay) and condition (massed, spaced).

| Choice test | Condition | Mean | CI |
| --- | --- | --- | --- |
| No-Delay | massed | 76.6 | [68.4 84.9] |
|  | spaced | 63.0 | [53.5 72.4] |
| Delay | massed | 75.4 | [66.3 84.5] |
|  | spaced | 76.1 | [66.1 86.2] |

| Accuracy change | Condition | Mean | CI |
| --- | --- | --- | --- |
| No-Delay | massed | -5.4 | [-14.5 3.7] |
|  | spaced | 5.8 | [-3.0 14.6] |
| Delay | massed | -13.8 | [-23.5 -4.1] |
|  | spaced | 18.2 | [8.2 28.2] |

**Figure 3.**
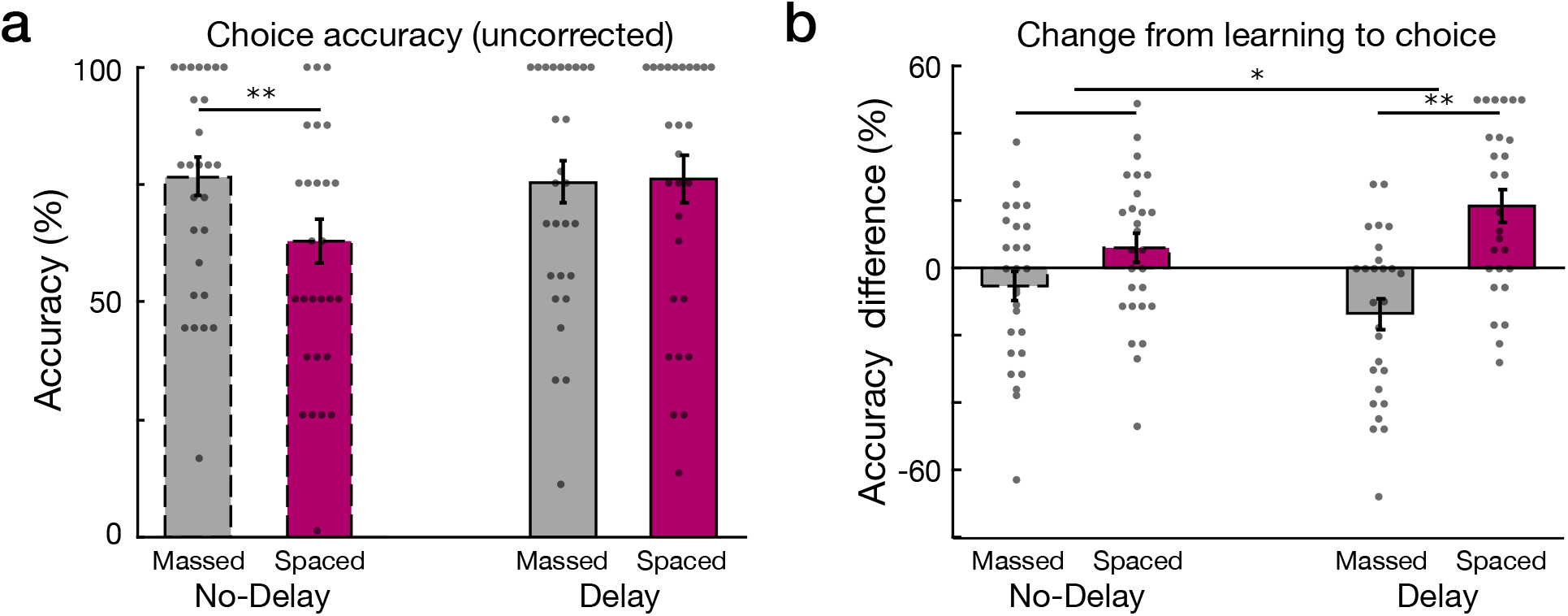
Choice test accuracy and adjusted preferences. (**a**) Uncorrected choice test phase accuracy for the No-Delay and Delay groups (massed condition in grey; spaced condition in magenta; outlines are dashed in the No-Delay group). (**b**) Change from the end of learning to the choice test phase. The interaction between delay group and spacing condition was significant. In the Delay group, the relative performance change was significantly different, reflecting a decrease in preference strength for massed-trained stimuli and an increase in preference strength for spaced-trained stimuli. Dots represent individual participants; outlines are dashed in the No-Delay group. (*p < 0.05; **p < 0.01; for underlying data, see https://osf.io/7wa3c/ and https://osf.io/38vzr/)

Next, we examined the critical question of whether a delay to test affected the maintenance of memory from the end of the learning phase to the choice test phase. We refer to this difference measure as change in preference strength. We found a significant interaction between delay group and spacing condition in post-learning preference (F_(2,104)_ = 5.15, p = 0.025; η_p_^2^ = 0.047; **Fig. 3b**). In planned comparisons, we found that in the No-Delay group, the difference in preference in the massed-trained condition numerically decreased while preferences in the spaced-trained condition numerically increased, although this interaction was not significant (t_(26)_ = −1.83, CI [−23.7 1.4]; p = 0.078; TOST p = 0.135; **Fig. 3b**). In the Delay group, the change in relative performance between the end of learning and test revealed a significant difference between spacing conditions (t_(26)_ = −5.07, CI [−45.0 −19.0]; p < 0.001; Fig. 3b).

The test phase choices employed a different measure than the learning phase responses, making it difficult to attribute preference change effects to either the massed or the spaced condition separately. However, in the No-Delay group, the numerical shift was similar and opposing for the massed and spaced conditions (massed: −5.4% CI [−14.5 3.7]; t_(26)_ = −1.22, p = 0.94, corrected for multiple comparisons; spaced: 5.8% CI [−3.0 14.6]; t_(26)_ = 1.36, p = 0.75, corrected). In the Delay group, the effects were in the expected direction (massed: −13.8% CI [−23.5 −4.1]; t_(26)_ = −2.92, p = 0.028, corrected; spaced: 18.2% CI [8.2 28.2]; t_(26)_ = 3.73, p = 0.0038, corrected).

These results indicate that rest has a detrimental effect on learned values for massed-trained stimuli, but a positive effect on learned values for spaced-trained stimuli. Overall, the decrease in massed relative to spaced condition preference strength supports a role for a working memory mechanism during learning that fails to maintain massed-trained value associations. Meanwhile, the relative increase in spaced condition preference strength supports the role of a separate mechanism that may increase the strength of value associations during rest.

### Post-learning reward ratings

Preceding the primary choice test measure, discussed above, we also collected a supplemental reward association rating for each stimulus. Participants rated each stimulus on a graded scale, anchored by 0% probability of reward and 100% probability of reward. Across groups, reward association ratings were significantly higher for reward- compared to loss-associated stimuli across spacing conditions (uncorrected ratings, effect of valence F_(3,208)_ = 110.16, p < 0.0001; η_p_^2^ = 0.346; all interaction p-values > 0.77; **Table 3**). Overall reward ratings were lower in the spaced condition than the massed condition (F_(3,208)_ = 8.90, p = 0.0032; η_p_^2^ = 0.041). We then tested the effect of delay on ratings, adjusted for performance at the end of learning. We found no significant interaction between delay group and spacing condition (corrected ratings F_(2,104)_ = 1.46, p = 0.23; η_p_^2^ = 0.014), unlike the results for the choice test. To further investigate this apparent difference between the choice test and the reward ratings results, we examined other effects of delay on reward ratings and the relationship between ratings and choices.

**Table 3.**
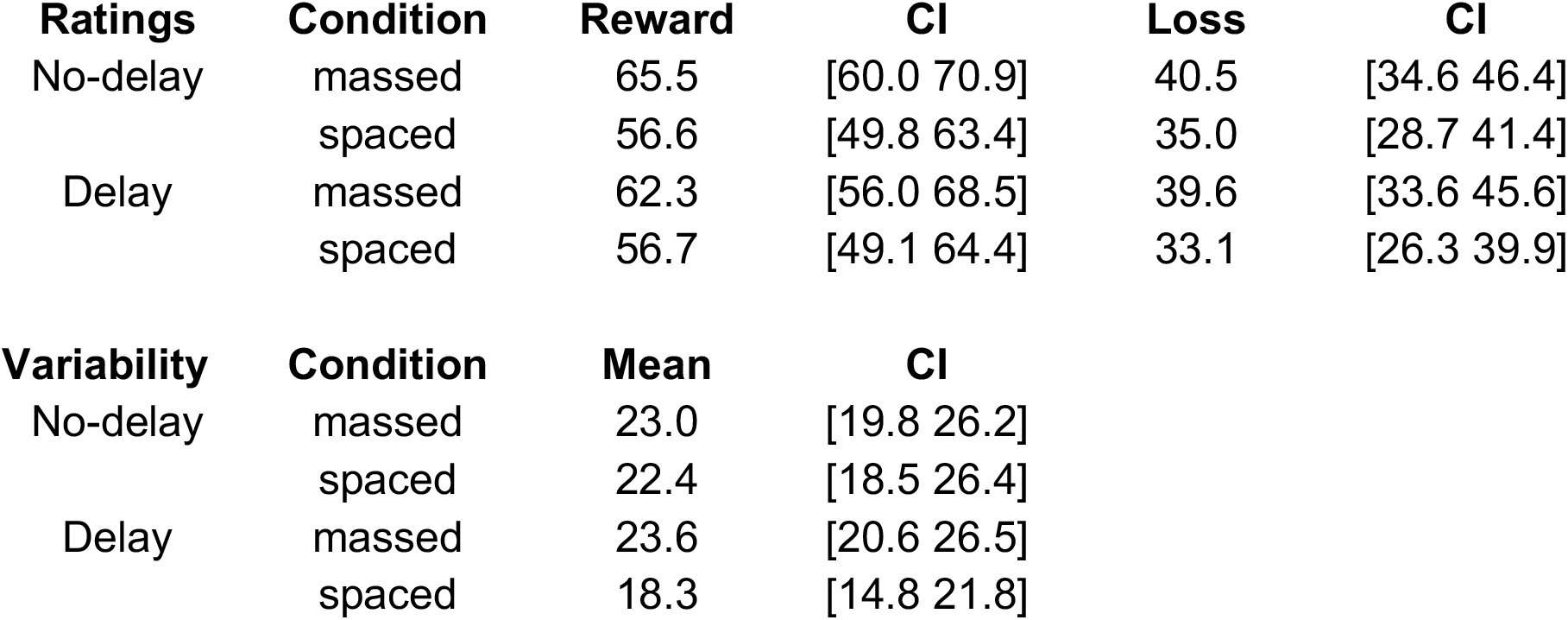
Post-learning reward ratings (top) for reward- and loss-associated stimuli, and ratings variability (bottom). Variability was defined as the mean of within-participants standard deviation (computed for reward and loss separately prior to averaging).

| **Ratings** | **Condition** | **Reward** | **CI** | **Loss** | **CI** |
| --- | --- | --- | --- | --- | --- |
| No-delay | massed | 65.5 | [60.0 70.9] | 40.5 | [34.6 46.4] |
|  | spaced | 56.6 | [49.8 63.4] | 35.0 | [28.7 41.4] |
| Delay | massed | 62.3 | [56.0 68.5] | 39.6 | [33.6 45.6] |
|  | spaced | 56.7 | [49.1 64.4] | 33.1 | [26.3 39.9] |

| **Variability** | **Condition** | **Mean** | **CI** |
| --- | --- | --- | --- |
| No-delay | massed | 23.0 | [19.8 26.2] |
|  | spaced | 22.4 | [18.5 26.4] |
| Delay | massed | 23.6 | [20.6 26.5] |
|  | spaced | 18.3 | [14.8 21.8] |

First, we examined whether the delay to test led to a relative increase in the variability of the reward ratings for massed- versus spaced-trained stimuli. We found that ratings variability for spaced versus massed stimuli was numerically but non-significantly lower after a delay (interaction F_(2,104)_ = 1.96, p = 0.164; η_p_^2^ = 0.019; **Table 3**; variability was computed as the average across the standard deviations for reward- and loss-associated stimuli separately). In planned comparisons, we found that in the No-Delay group there was no difference in ratings variability between massed- and spaced-trained stimuli (t_(26)_ = 0.26, CI [−3.9 5.0]; p = 0.797). In the Delay group, however, we found that ratings variability for spaced-trained stimuli was lower than variability for massed-trained stimuli (t_(26)_ = −2.44, CI [−9.7 −0.8]; p = 0.022).

Next, we examined how strongly reward ratings were linked to choices, as higher variability may decrease this relationship. For each participant, we estimated a model relating per-stimulus ratings to mean choice preference for a stimulus, separately for massed and spaced-trained stimuli. As expected, we found a positive relationship overall between post-learning ratings and choice preference (t_(53)_ = 15.94, p < 0.0001). Comparing the relationship between ratings and choices across the delay groups, we found that a delay before the test led to a significant interaction, reflecting an increase in the relationship between ratings and choices in the spaced versus the massed condition (interaction F_(2,104)_ = 4.46, p = 0.037; η_p_^2^ = 0.041). In planned comparisons, we found that the ratings-choice relationship in the No-Delay group did not differ based on spacing condition (t_(26)_ = 1.08, CI [−0.001 0.004]; p = 0.29; TOST p = 0.036), while the ratings-choice relationship in the Delay group was significantly weaker in the massed compared to the spaced condition (t_(26)_ = −2.37, CI [−0.007 −0.001]; p = 0.026). Finally, exploratory analyses suggested that the difference in the ratings-choice relationship between the No-Delay and Delay groups was driven by the spaced condition (massed t_(52)_ = 0.40, p = 1, corrected; spaced t_(52)_ = −2.57, p = 0.026, corrected).

Thus, while preferences measured via incentive-compatible choices were our primary outcome measure, unlike the results for choices, we found no significant interaction of delay group and spacing condition on post-learning reward ratings. However, we found that after a rest delay, ratings for spaced-trained stimuli were relatively better predictors of choices. This pattern is consistent with a decay in the fidelity of massed-trained value associations and an increase in the fidelity of spaced-trained value associations. We suggest that these underlying patterns in the reward rating data are consistent with opposing effects of a rest delay on the variability of learned values and choice accuracy for massed-versus spaced-trained stimuli.

## Discussion

We examined the effect of massed versus spaced learning of reward and loss associations and the effect of post-learning rest on memory for value associations. The close repetition of trials in our “massed” condition was designed to be similar to many feedback-based learning designs used in human research (e.g. Daw et al., 2011; Wimmer et al., 2012; Wimmer et al., 2014). During learning, repetitions for massed stimuli were presented sequentially with occasional interruptions for the presentation of “spaced” stimuli. Across multiple measures and two experimental groups, we found that short-term working memory resources supported massed performance during learning. First, we found that massed training led to better overall performance during the learning phase. Second, brief interruptions of massed training were immediately followed by performance decreases. Third, an independent measure of working memory capacity was related to massed but not spaced learning performance. These results build on previous findings that reward learning in humans with closely-spaced repetitions is supported by an interaction between working memory and reinforcement learning mechanisms (Collins and Frank, 2012; van de Vijver et al., 2015; Collins and Frank, 2018; Wimmer et al., 2018; van de Vijver and Ligneul, 2019). Finally, we found an effect of a brief post-learning delay: when tested with no delay, relative preference strength was similar to performance at the end of learning. However, when tested after a brief delay, relative preference strength for massed-trained stimuli decreased while preference strength for spaced-trained stimuli increased. Thus, working memory may aid short-term performance while at the same time negatively interacting with reinforcement learning processes responsible for longer-term maintenance (Collins, 2018).

### Working memory and learning phase performance

Our results during the learning phase provide new support for an important role of short-term memory in typical reward-based and reinforcement learning paradigms. Recently, in conditions resembling massed learning, we found a positive correlation between working memory and learning performance (Wimmer et al., 2018). Previous work supporting a negative interaction between episodic memory encoding and reinforcement learning suggested that this interaction may be due to competition over short-term memory processes and attention (Wimmer et al., 2014). Building on this work, the current paradigm provides both individual difference and trial-by-trial measures of the interaction between working memory and reward-based or reinforcement learning in the same task. Further, our novel implementation of a concurrent dual-task during reward learning kept massed-trained performance below ceiling, which may help to reveal links between learning and working memory.

A closely related line of research on working memory and learning has indirectly investigated the effect of within-session spacing (Collins and Frank, 2012; Collins et al., 2014; Collins et al., 2017; Collins, 2018; Collins and Frank, 2018). In the paradigm used by Collins et al., participants learn which of 3 response options are reinforced for a given stimulus, while across separate blocks, the number of concurrently learned stimuli (“load”) is systematically manipulated. Notably, by changing the number of stimuli concurrently presented within a block, this task also manipulates the spacing between repetitions of a given stimulus. While reinforcement learning models that only acquire stimulus-action values (“model-free” RL) predict that the number of stimuli in a block should have no effect on performance, strikingly, Collins et al. find that ongoing learning performance is significantly decreased as more stimuli are included in the learning block. This performance decrease with increasing load was successfully modeled using a reinforcement learning model that included a fixed-capacity working memory module capable of maintaining stimulus-response mappings (see also van de Vijver and Ligneul, 2019).

More recently, the effect of learning phase load on later choices was investigated by Collins et al. (2017; 2018), with several similarities to the current paradigm’s use of separate learning and choice test phases. In these studies, the test phase followed a distractor task at a very similar delay to that used in our Delay group. The authors report relatively worse test phase accuracy for stimuli learned in low set-size blocks (similar to the current massed condition), suggesting that short-term working memory supports learning and then fails to support later test accuracy (Collins et al., 2017; Collins, 2018).

The current work extends these findings in several ways. We combine conditions favoring short-term over longer-term learning into the same relatively short learning session (see also van de Vijver and Ligneul, 2019). We find that an independently collected measure of working memory capacity correlates with massed (but not spaced) learning performance (van de Vijver et al., 2015; Wimmer and Poldrack, 2018). When tested after no delay, we find accuracy benefits for massed-trained stimuli that align with previous results (Collins, 2018). However, by manipulating a post-learning delay across groups, we show that relative performance for massed-trained stimuli decays across a brief delay, such that preferences were weaker for massed-trained stimuli but stronger for spaced-trained stimuli.

We found two timescales of decay in massed performance: an immediate decay due to trial-by-trial interruption and a slower preference decay across a post-learning delay. These different effects suggest extensions or modifications to the single probabilistic working memory model proposed previously (Collins and Frank, 2012). Our current results and related findings (Wimmer and Poldrack, 2018) indicate that an alternative conceptualization of working memory may be necessary. Instead of a single working memory store with a single rate of forgetting, graded or multiple memory stores that support very short-term through to longer-term storage may better capture the full range of observed behavior (Eldar et al., 2018). Further, our results support the perspective that working memory representations are not binary but are graded in fidelity (Ma et al., 2014).

### Effect of post-learning delay period

Critically, we found an effect of rest delay on memory for value associations, where relative performance decreased for massed-trained stimuli and increased for spaced-trained stimuli. Our results suggest that a post-learning delay period may both weaken preferences for massed-trained value associations and strengthen preferences for spaced-trained value associations (Schmidt and Bjork, 1992; Taylor and Rohrer, 2010). However, we cannot directly attribute these changes to effects of either the massed or spaced training alone. The direct comparisons of performance from the No-Delay to the Delay groups for massed- or spaced-trained stimuli were not significant, although our results indicate that we cannot rule out the presence of a medium-sized effect in each condition (especially for spaced-trained stimuli). For spaced-trained items, the change in performance at test cannot be explained as a simple effect of impaired performance during learning, as learning phase performance was similar in both the No-Delay and Delay groups. One possibility is that value associations that are weakly encoded during the learning phase are “sharpened” during the post-learning delay period. This interpretation is further supported by the significant decrease in post-learning reward rating variability after a delay.

Previous research on the effects of spacing has shown that performance is relatively impaired by spaced training while later retention is improved (Schmidt and Bjork, 1992; Taylor and Rohrer, 2010; Soderstrom and Bjork, 2015), similar to our learning phase results. Additionally, research in this area has also reported a positive effect of a post-learning delay on test performance for verbal memory and motor learning (Lee and Genovese, 1988; Donovan and Radosevich, 1999; Janiszewski et al., 2003; Cepeda et al., 2006; McCabe, 2008). This area of research makes a distinction between short-term retention versus later evidence of learning (Schmidt and Bjork, 1992; Taylor and Rohrer, 2010; Soderstrom and Bjork, 2015), a conceptualization which can also map to our findings on massed versus spaced training. From this perspective, many reward and reinforcement learning studies in humans could be considered to be studying the mechanisms supporting short-term retention instead of mechanisms supporting lasting learned values or preferences.

Positive effects of a delay period could arise from several not mutually exclusive mechanisms. Based on recent neuroscience research, one mechanism could be spontaneous reactivation of spaced-trained associations after learning (Gomperts et al., 2015; Gruber et al., 2016; Olafsdottir et al., 2018), which can be implemented computationally in the DYNA reinforcement learning model (Sutton, 1990). Such a mechanism has been proposed to support positive effects of rest on learning in a multi-step associative task (Gershman et al., 2014), and may also relate to implicit covert retrieval processes during and after learning (McCabe, 2008). Second, spacing may also lead to more lasting learning due to the unexpected nature of spaced item appearances during training combined with the requirement for memory retrieval (Bouton and Moody, 2004). Relative novelty has been demonstrated to interact with reward processing, leading to increased activity in the hippocampus as well as the striatum (Guitart-Masip et al., 2010; Bunzeck et al., 2011; Zaehle et al., 2013). This effect may be driven by a hippocampal-VTA loop, supporting an increase in the firing rate of dopamine neurons in response to rewards for spaced stimuli (Lisman and Grace, 2005; Goto and Grace, 2008). Finally, spacing of learning repetitions may also allow for relatively short-term synaptic plasticity mechanisms to iteratively build stronger associations (Reynolds et al., 2001), potentially aided by dopamine release during learning (Grogan et al., 2017).

It is possible that a longer delay before testing would reveal more robust changes in performance at test. Supporting this view, two previous studies have found effects of spacing after at least one week, allowing for sleep-related consolidation processes (Wimmer et al., 2018; van de Vijver and Ligneul, 2019); here, training was either spaced across days (Wimmer et al., 2018) or within a single session (van de Vijver and Ligneul, 2019). In the latter study, when a test was given immediately after learning, the authors found no change in performance from the end of learning (van de Vijver and Ligneul, 2019), aligning with the choice test results in our No-Delay group. By using a brief delay, however, our paradigm demonstrates that effects of spacing on the maintenance of learning can be successfully studied in just a single session.

## Conclusion

In summary, we found that reward-based learning with typical massed presentation of stimuli shows a strong dependence on short-term working memory resources across multiple measures. After learning, a delay period revealed that preferences for spaced- versus massed-trained were better maintained. These and other related findings indicate that studies of human reward-based learning that employ common condensed learning designs may be a suboptimal way to measure individual differences in the mechanism supporting lasting reward-based or reinforcement learning (Collins and Frank, 2012; Collins et al., 2014; Collins, 2018; Wimmer et al., 2018; van de Vijver and Ligneul, 2019). These results have implications for the design and interpretation of research on reward-based learning in learning deficits in psychiatric disorders (Maia and Frank, 2011; Montague et al., 2012; Whitton et al., 2015; Huys et al., 2016; Moutoussis et al., 2016).

To better isolate reward-based learning mechanisms in humans, in particular the role of the striatal dopamine system, our results support the use of experimental designs with increased spacing between training repetitions. Spaced designs can also increase our ability to understand behavior outside the lab, where learning repetitions are often spread over periods of time longer than several seconds. Importantly, spacing training across days can produce value associations that are resistant to forgetting (Kim et al., 2015; Wimmer et al., 2018; van de Vijver and Ligneul, 2019), reminiscent of lasting habits. The increased maintenance of spaced associations may be particularly relevant for understanding maladaptive value associations, such as those found in addiction.

## Acknowledgments

The authors thank Jamie Li for assistance in data collection and helpful discussions. Research was supported by NIH R01AG041653 [RP] and a research fellowship from the Deutsche Forschungsgemeinschaft [GEW].

## Open Practices Statement

The full behavioral dataset is available at OSF (https://osf.io/xu48j/). None of the experiments were preregistered.

## Supplementary Results

### Choices across massed and spaced conditions

We also collected choices between massed- and spaced-trained stimuli, comparing reward- versus reward-associated stimuli and loss- versus loss-associated stimuli. In the No-Delay group, for reward-associated stimuli, we found that participants preferred massed-trained stimuli over the spaced-trained stimuli (massed versus spaced: 64.2% CI [56.4 71.9]; t_(26)_ = 3.767, p < 0.001), consistent with the strength of working memory-supported values for massed-trained stimuli at no delay. For loss-associated stimuli, we found no difference in preference (51.5 % CI [39.0 58.0]; t_(26)_ = 0.334, p = 0.74; TOST p = 0.007). Reflecting the shift in relative preference across the delay, in the Delay group we found that participants showed no preference for massed- or spaced-trained stimuli (reward-associated massed versus spaced: 52.5% CI [43.8 61.1]; t_(26)_ = 0.588, p = 0.56; TOST p = 0.007; loss-associated: 49.7% CI [40.1 59.3]; t_(26)_ = −0.066, p = 0.95; TOST p = 0.004).

Comparing performance in these mixed choices across the No-Delay and Delay groups, we found that for reward-associated stimuli, preferences for massed- versus spaced-trained stimuli were significantly lower in the Delay versus No-Delay group (reward difference: −11.7% CI [−23.0 −0.4]; t_(52)_ = −2.079; p = 0.043; loss difference, −1.9% CI [−15.0 11.3]; t_(52)_ = −0.28; p = 0.779; TOST p = 0.011). This relative decrease in value for massed and increase in value for spaced reward-associated stimuli after rest parallels the changes in performance from learning to test.

